# Mobility of the human foot’s medial arch enables upright bipedal locomotion

**DOI:** 10.1101/2022.09.13.507861

**Authors:** Lauren Welte, Nicholas B Holowka, Luke A Kelly, Toni Arndt, Michael J Rainbow

## Abstract

Developing the ability to habitually walk and run upright on two feet is one of the most significant transformations to have occurred in human evolution. Many musculoskeletal adaptations enabled bipedal locomotion, including dramatic structural changes to the foot and, in particular, the evolution of an elevated medial arch (H. Elftman and Manter, 1935). The foot’s arched structure has previously been assumed to play a central role in directly propelling the centre of mass forward and upward through leverage about the toes (Herbert Elftman and Manter, 1935) and a spring-like energy recoil (Hicks, 1955). Paradoxically, these roles seemingly require either arch rigidity (for the former) or mobility (for the latter). However, it is unclear whether or how the mobility and height of the medial arch support its propulsive lever function. Here we show, using high-speed biplanar x-ray, that regardless of intraspecific differences in medial arch height, arch recoil enables a longer contact time and favourable propulsive conditions for walking upright on an extended leg. This mechanism presumably helped drive the evolution of the longitudinal arch after our last common ancestor with chimpanzees, who lack this mobility during push-off. We discovered that the previously overlooked navicular-medial cuneiform joint is primarily responsible for this mobility in human arches, suggesting that future morphological investigations of this joint will provide new interpretations of the fossil record. Our work further suggests that enabling the mobility of the longitudinal arch in footwear and surgical interventions is critical for maintaining the ankle’s natural propulsive ability.

## Introduction

The foot experienced strong selective pressures during human evolution. Features unique to the human foot, such as a pronounced medial arch, have been proposed to play a key role in the evolution of habitual bipedalism (H. Elftman and Manter, 1935). The presence of a high medial arch in fossil hominins has been argued to represent an adaptation for both the foot’s rigidity in push-off (Herbert Elftman and Manter, 1935; Sarrafian, 1987; Susman, 1983) and its mobility-enabled spring-like function (Hicks, 1955; Holowka and Lieberman, 2018; McNutt et al., 2018). The rationale for these ideas is that, similar to ancient architecture, a curved longitudinal arch provides a nearly rigid lever for push-off (Morton, 1924). Simultaneously, the mobility of various foot joints enables the spring-like recoil of the arch-spanning tissues (Kelly et al., 2014; Ker et al., 1987; Stearne et al., 2016). However, it is unclear how the coupled mobility-rigidity functions work in tandem – or contradictorily – to enable propulsion and if they are linked to the evolution of upright bipedalism.

Relative to our arboreal primate relatives, hominins are widely considered to have rigid feet to enable efficient leverage in propulsion (H. Elftman and Manter, 1935; Griffin et al., 2010). However, human feet are mobile-and not rigid-undergoing more medial arch recoil than chimpanzee feet in walking propulsion (Holowka et al., 2017a). In addition, many primates (including humans) take advantage of mobility-enabled energy storage mechanisms in their arch-spanning tissues (Bennett et al., 1989; Vereecke and Aerts, 2008). The mobility of human and non-human primates’ medial arches may therefore influence foot leverage in bipedal locomotion. The distinct locomotor difference between the hominin and panin lineages is the position of the medial arch immediately following heel lift. Non-human primates overwhelmingly experience a midfoot break (D’Août et al., 2002; DeSilva, 2010; Herbert Elftman and Manter, 1935; Griffin et al., 2010; Vereecke et al., 2003). The hindfoot lifts relative to the ground-contacting metatarsals, forming at times a “reverse-arch” in which the midfoot is below the plane connecting the heel and the toes (Bennett et al., 1989). The fulcrum of the foot-lever becomes the midfoot instead of the metatarsophalangeal joints (Griffin et al., 2010), shortening the foot-lever and reducing the levering advantage. This arch mobility may provide advantages for climbing (Bojsen-Møller, 1979; Herbert Elftman and Manter, 1935; Holowka et al., 2017b) but imposes challenges in pushing off into the next step when walking bipedally. In contrast, humans’ robust plantar fascia and plantar ligaments, as well as the elevated arch structure, contribute to the stability of the longitudinal arches (Bojsen-Møller, 1979; DeSilva, 2010; Sichting et al., 2020). At heel lift, they keep the midfoot above the plane connecting the heel and the toes. Some humans experience a slight midfoot break, but not to the extent of other primates (DeSilva et al., 2015; Greiner and Ball, 2014). As a result, the fulcrum of the human foot-lever is primarily the metatarsophalangeal joints (Griffin et al., 2010; Hicks, 1954), enabling humans to take advantage of additional leverage relative to primates with a midfoot break. While non-human primate arch mobility influences the fulcrum position of the foot-lever in bipedal locomotion, it remains unclear how human arch recoil influences its function as a lever. Understanding this mechanism may further elucidate our evolutionary divergence from other primates.

Human foot arch recoil (Holowka et al., 2017a; Holowka and Lieberman, 2018; Ker et al., 1987; Stearne et al., 2016; Vereecke and Aerts, 2008) and rigid foot levering (Kondo et al., 2021; Susman, 1983) are both thought to assist with centre of mass propulsion in locomotion (Cavagna et al., 1977) (Figure 1a). In late stance, arch-spanning tissue recoil shortens and lifts the arch, theoretically helping to move the centre of mass forwards and upwards (Kelly et al., 2014; Ker et al., 1987; Stearne et al., 2016). Concurrently, the calf muscles’ substantial power generation lifts the ankle (Farris and Sawicki, 2012; Neptune et al., 2001); however, the foot must be sufficiently stiff to enable the rising ankle’s contribution to centre of mass propulsion (Cavagna et al., 1977). In concept, if the foot is perfectly rigid (i.e., with an arch that does not recoil), it can lever efficiently off the ground as it rotates around the metatarsal-phalangeal joints (Griffin et al., 2010). In reality, the foot is not completely rigid but has features that increase its intrinsic stiffness and limit midfoot break. The plantar muscles (Farris et al., 2019; Kelly et al., 2014), ligaments (Bates et al., 2013; Bojsen-Møller, 1979; Ker et al., 1987; Lovejoy et al., 2009), and fascia (Ker et al., 1987; Sichting et al., 2020) work in tandem with the shape and posture of the bones in the medial (Hicks, 1955; H. Elftman and Manter, 1935; Bojsen-Møller, 1979), lateral(McNutt et al., 2018), and transverse arches (Venkadesan et al., 2020) to increase the intrinsic stiffness that enables foot leverage. Specifically, the static height of the medial arch is thought to appreciably influence the arch’s intrinsic stiffness, despite the generally weak relationship between arch height and arch vertical mobility (Cornwall and McPoil, 2011; Zifchock et al., 2006). A higher and stiffer arch would presumably enable more efficient propulsion by improving the arch’s rigid-levering behaviour. Thus, we tested the contributions of medial arch height and mobility to centre of mass propulsion. We predicted that a higher unloaded medial arch would propel the centre of mass more than a flatter arch. We also hypothesised that arch recoil would increase the height and forward progression of the centre of mass above the contributions of the levering motion.

**Figure 1.**
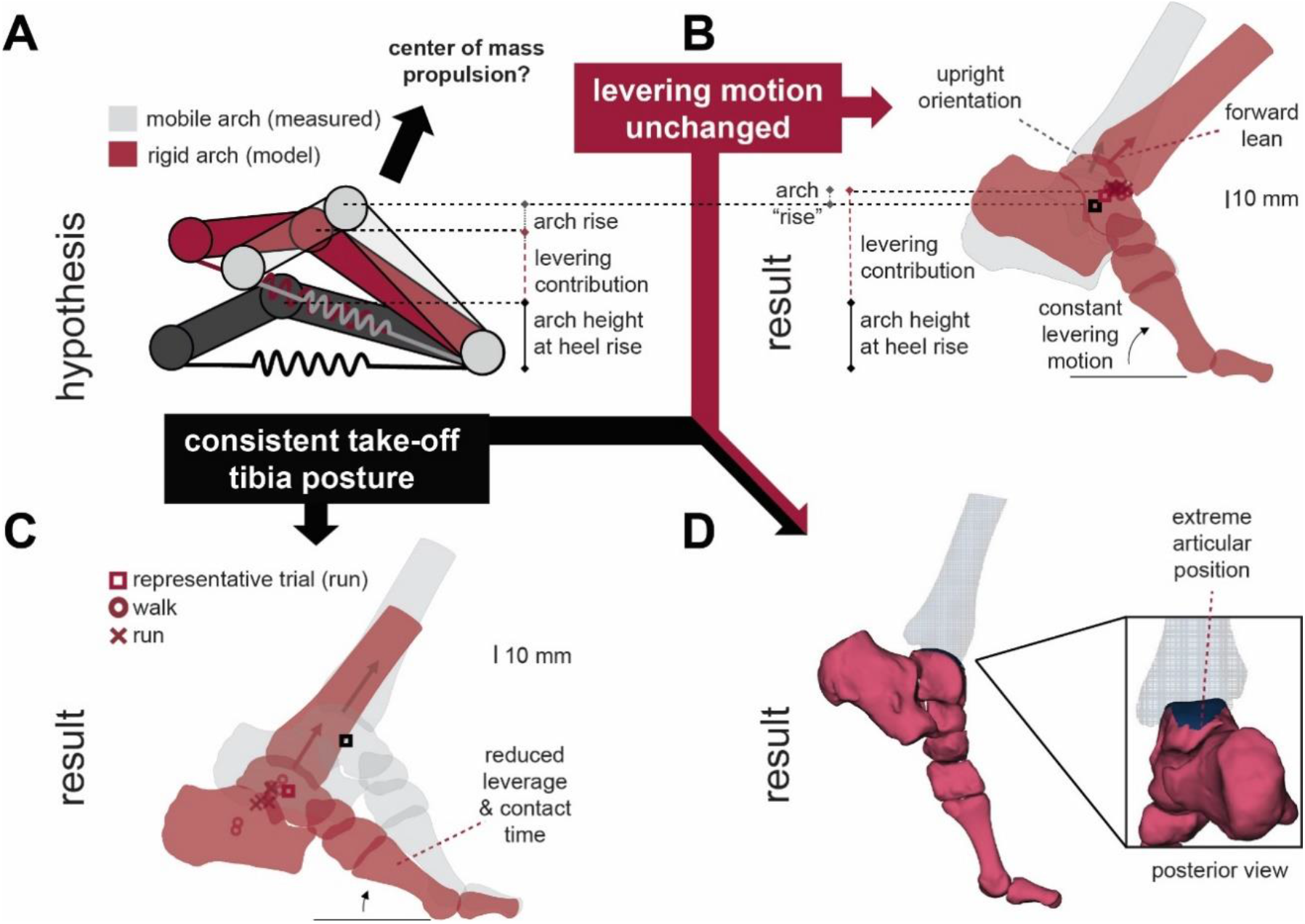
The contributions of the rigid-lever and mobile arch to the propulsion of the centre of mass. (a) Conceptual diagram of the foot’s hypothesised contributions to centre of mass propulsion. The ankle is lifted through a combined rigid-levering (red) about the metatarsal heads and the combined levering and recoil of the medial arch (grey). For the high-speed x-ray measured motion (grey) and the rigid-lever model (red), the position of the talar centroid, as a proxy for centre of mass propulsion, are shown at three different push-off conditions: (b) with the same metatarsal levering motion as the measured bone motion, which results in substantial forward lean; (c) with the same posture of the tibia at push-off, which would reduce levering and contact time with the ground, and (d) with both the same levering and tibia posture, which requires an extreme articular position and more rapid ankle plantarflexion. Note that the posterior view shows the tibia moved past the articular surface of the talus.

## Results & Discussion

To evaluate the individual contributions to centre of mass propulsion of the mobile, rising arch and the levering of a rigid foot, we used *in vivo* three-dimensional measurements of the foot skeleton during locomotion. Seven participants walked and ran at self-selected speeds (walking 1.6 ± 0.1 m/s, running 3.0 ± 0.4 m/s) over a raised walkway while high-speed biplanar x-ray images were captured (walking 125 Hz, running 250 Hz). A semi-automatic shape matching process (Akhbari et al., 2019; Miranda et al., 2011) animated three-dimensional models of the bones forming the ankle joint complex and medial arch (first metatarsal, medial cuneiform, navicular, talus, calcaneus, and tibia), as well as the first proximal phalanx. Starting just before heel lift, we modelled a rigid arch-lever by mathematically fixing the bones in the medial arch to the first metatarsal (the fulcrum of the foot-lever) while allowing motion at the ankle. This approach decoupled the foot’s levering behaviour from arch mobility. Talar centroid height and forward progression were a proxy for centre of mass propulsion. Contrary to our expectations, the recoiling motion of the arch did not directly propel the centre of mass forwards or upwards above the foot’s levering (Figure 1b). Talar centroid height for the rigid arch was 5.7 ± 2.4 % (p < 0.01, r = 1) higher and 11.0 ± 3.2 % (p < 0.01, r = 1) more forward (superior: 70.2 ± 13.3 mm, anterior: 71.8 ± 20.3 mm) than when the mobility of the arch was maintained (superior: 66.5 ± 13.1 mm, anterior: 64.5 ± 17.5 mm).

This surprising result may lead us to falsely conclude that a rigid foot would be more effective at propelling the centre of mass than a mobile arch; however, our data show that arch recoil determined the upright posture of the talus. The recoiling medial arch causes the talus to rotate backwards, curling under the tibia during ankle plantarflexion (see Supplemental Video 1). The tibiotalar articular surface of the talus thus faces superiorly at push-off but is lower than the modelled rigid arch. This change in orientation and position of the talus would have important ramifications for the tibia’s posture, likely influencing motion at the knee and hip. For example, our model shows that when tibiotalar motion and levering motion are unchanged, the arch as a perfect rigid lever results in substantial anterior lean of the tibia compared to the recoiling arch condition (Figure 1b, Supplemental Video 1). Accommodating the tibial lean produced by a rigid arch requires push-off with a flexed knee, which would likely reduce the efficiency of human bipedal movement (Waters and Mulroy, 1999). Alternatively, to push off with a typical upright tibia posture (Figure 1c), our model shows that the rigid arch would have 29.8 ± 9.0% (0.028 ± 0.008 s) less propulsive ground contact time and impulse would be substantially reduced (Figure 1c). As a result, propulsion would likely require larger muscle forces and higher contraction velocities. Alternatively, if we allow the same levering motion and tibia lean at push-off with a rigid foot as we observe with a mobile foot, the tibiotalar joint would achieve an extreme articular position with less overlap between joint surfaces (Figure 1d). Additional ankle plantarflexion would be required for the same time period, increasing ankle plantarflexion velocity and potentially forcing the calf muscles to generate propulsive power under unfavourable contractile conditions (Carrier et al., 1994).

We aimed to understand which of our modelled cases (Figure 1) is most similar to how humans behave during locomotion and if the medial arch height contributes to foot-lever-driven propulsion. Using multiple linear regression, we investigated the influence of participant arch height and mobility on talar propulsion and ankle plantarflexion. Our seven subject sample was selected to span a range of arch heights from 20 initially enrolled participants. Therefore, our model contains a distribution of flat to high static arch positions (approximate seated arch height index of 0.33±0.03, range 0.29-0.39; healthy human average (Butler et al., 2008) is 0.36±0.03). We used predictor variables of arch recoil range of motion, unloaded static arch angle, and a categorical variable for walking and running. Our multiple linear regression model showed that arch recoil range of motion was a better predictor of the magnitude of anterior and superior displacement of the talar centroid (R^2^ = 0.82, p < 0.01, 95%CI: [-5.9,-3.2]) than unloaded static arch angle (p = 0.79, 95%CI: [-0.7,0.3]). As arch recoil increases, talus propulsion increases. Our rigid arch model suggested that arch recoil does not directly lift the centre of mass over and above the lever action of the foot. However, it seems that in vivo, participants take advantage of additional arch recoil, which leads to an upright-facing talar articular surface and enables propulsion from a higher ankle position (consistent with the model in Figure 1c). Using the same independent variables, we found that arch recoil range of motion better predicted ankle plantarflexion range of motion in propulsion (R^2^ = 0.92, p < 0.01, 95% confidence interval on slope (95%CI) : [0.4,1.1]) than unloaded static arch angle (p = 0.79, 95%CI [-0.2,0.1]). As a person has a more mobile medial arch, the superior talar surface is more upright, increasing the tibia’s available range of motion while remaining relatively vertical. In other words, consistent with our rigid arch model (Figure 1d), the range of tibiotalar motion that keeps the tibia upright is limited by the superior talar surface’s upright orientation and geometry. Arch and ankle plantarflexion range of motion were also significantly larger in running than in walking, suggesting that arch mobility may play a more critical role in maintaining an upright tibia posture in running. This is further emphasised by the larger effect of a rigid arch compared to the mobile arch on global tibia posture in running compared to walking (Figure 2ab). Since the postural advantages of arch mobility are present in walking but play a larger role in running, it is possible that evolutionary pressure for a mobile medial arch was amplified as humans began to run(Bramble and Lieberman, 2004). Our results further imply that humans have an optimal tibia orientation for propulsion, which may be limited by the arch’s ability to recoil. Thus, arch mobility is coupled to propulsive ankle mechanics, while intraspecific static arch height is not.

**Figure 2.**
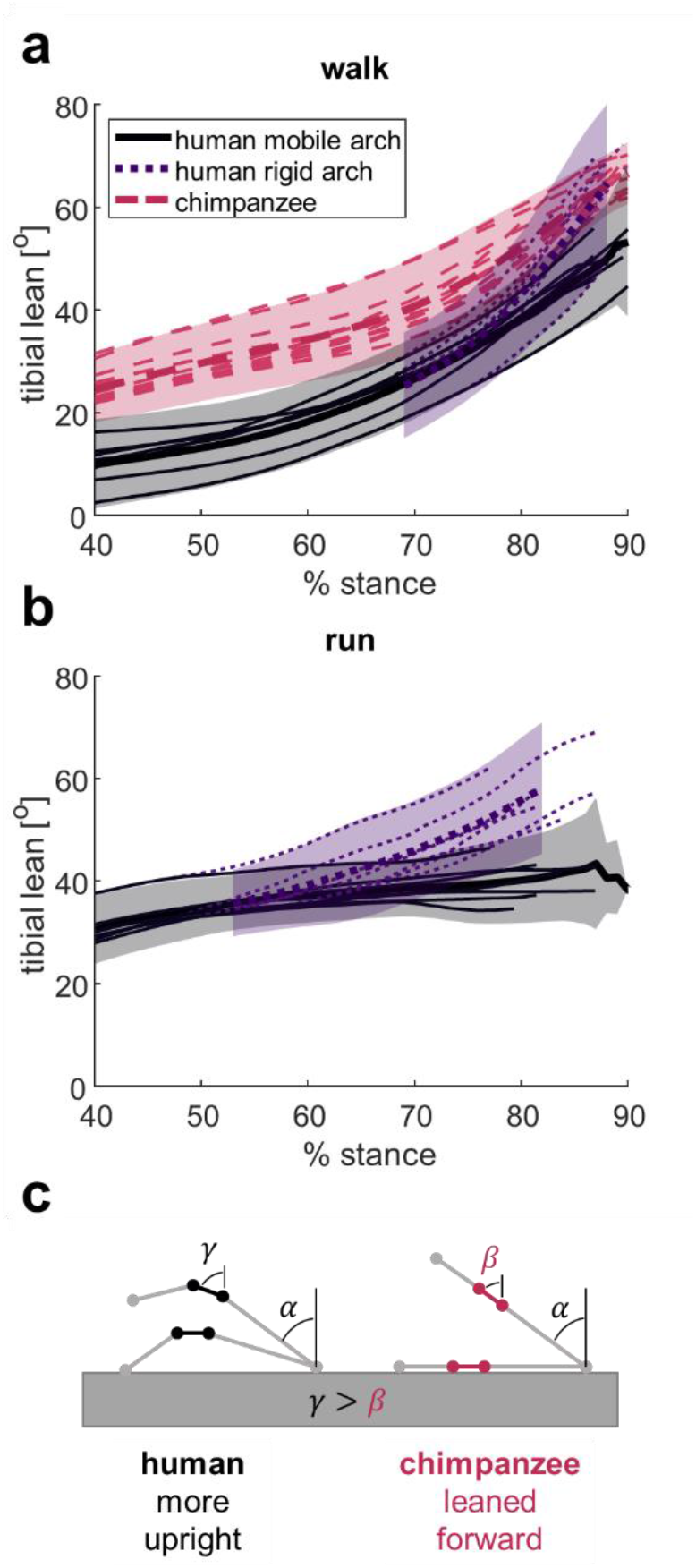
Tibia lean as measured in human and chimpanzee locomotion. (a) The measured human mobile arch’s tibia orientation (black, solid line) is compared to the modelled rigid arch’s tibia (purple, dotted line) (N=7, 1 step/participant) and chimpanzee tibia orientation in bipedal walking (pink, dashed line) (N=3, 13 steps). At propulsion, the human rigid arch’s tibia is in the same position as the chimpanzee’s. Mean angles are shown in a thicker line, with ± 2 SD. (b) The human mobile arch (black, solid line) compared to the modelled rigid arch (purple, dotted line) in running. (c) The talus/midfoot line segment (black for the human rigid arch, pink for chimpanzee) at the same global metatarsal position in the levering process, without arch mobility. The talus/midfoot segment is naturally more superiorly oriented in the human arch. Human recoil further orients the talus upright, while the midfoot break in chimpanzees would further lean the talus forward.

While we showed that arch mobility enables an upright tibia posture, simply having an arched foot may help humans move bipedally compared to other primates. Humans locomote over an extended lower limb, thus requiring an upright tibia orientation compared to chimpanzees’ flexed limb walking posture (Figure 2a). We propose that humans have evolved a pronounced medial arch for two non-mutually exclusive reasons. First, the arch naturally orients the talus’ superior articular surface upright, such that even without arch recoil, it is more upright than in the non-arched feet of chimpanzees (Figure 2c). Secondly, the arch-spanning tissues in humans have a longer moment arm about the midfoot joints to produce more arch recoil than in chimpanzees. Thus, in addition to the natural upright orientation of the human talar superior surface, arch recoil further enables propulsion over a longer period while the tibia can remain upright. When chimpanzees experience a midfoot break, the talus and tarsal bones lean forward, which likely contributes to chimpanzees’ flexed-limbed posture in bipedal push-off. Due to their lack of arch, the midfoot has less capacity to recoil and re-orient the talus to be upright. These ideas are consistent with reduced midfoot recoil and reduced ground contact times in chimpanzees compared to humans (Holowka et al., 2017a; Sockol et al., 2007). Overall, a prerequisite for hominins to push off efficiently with a fully extended leg was the evolution of a structural arch to function in tandem with the recoiling arch.

The earliest fossil hominin with significantly preserved foot bones, *Ardipithecus ramidus*, lacked a medial arch (Lovejoy et al., 2009), indicating that our earliest ancestors likely utilised a primitive form of bipedalism that entailed walking on flexed limbs or with limited ankle plantarflexion during push-off. In support of the latter possibility, *A. ramidus* possessed a narrow posterior dorsal articular surface of its talus (DeSilva et al., 2019), suggesting limited weight-bearing capacity in plantarflexion. While current interpretations have focused on the presence of a medial arch for inferring efficient bipedal gait, our findings demonstrate that arch mobility, not arch height, enables upright locomotion. It is, therefore, possible that *A. ramidus* could still have possessed a mobile midfoot that allowed for some midfoot recoil and thus more upright bipedalism, even in the absence of an elevated medial arch. The australopithecines, which appeared slightly later in the fossil record, show the first evidence of a rudimentary medial arch alongside a wider posterior tibiotalar joint surface (DeSilva et al., 2019). As hypothesised in Figure 2c, the australopithecines could take further advantage of their midfoot mobility with the presence of an elevated medial arch, leading to morphological changes that enable additional ankle plantarflexion. These features coincide with a suite of other novel adaptations for bipedalism and are consistent with the evolution of a more human-like form of bipedalism on more extended legs (Pontzer, 2017). Later fossils and footprints attributed to *Homo* show evidence of more pronounced medial and transverse structural arches, likely supported by arch-spanning ligaments and muscles (Susman, 1983; Venkadesan et al., 2020). As a result, *Homo* could better take advantage of arch mobility and its effects on ankle posture and the contractile state of the triceps surae muscles in propulsion. The evolutionary scenario described above represents a hypothesis about the selective forces that drove the evolution of the medial arch, which requires further testing to establish the relationships between foot joint morphology and mobility.

The midfoot break, which has received much attention in previous studies of human foot evolution, is perceived to occur at the talonavicular, calcaneocuboid and cuboid-metatarsal joints(DeSilva, 2010; H. Elftman and Manter, 1935). Interestingly, we discovered that in propulsion in humans, medial arch mobility occurs primarily at the navicular-medial cuneiform joint (cuneonavicular joint) (Figure 3). While the cuneonavicular joint is known to be mobile (Arndt et al., 2007), our results indicate that it is primarily responsible for orienting the superior talar surface upright and enabling locomotion on extended limbs. In axial loading, gorillas and chimpanzees generally have less motion than humans between the navicular and first metatarsal (Negishi et al., 2021), which, based on our results here, we presume occurs at the cuneonavicular joint. Thus, our divergent evolutionary path from non-human primates toward obligate bipedalism is likely reflected in the morphology and mobility of the cuneonavicular joint. We suggest that future comparative and paleontological analyses should investigate changes in the morphology of the cuneonavicular joint to better understand the evolution of the medial arch.

**Figure 3.**
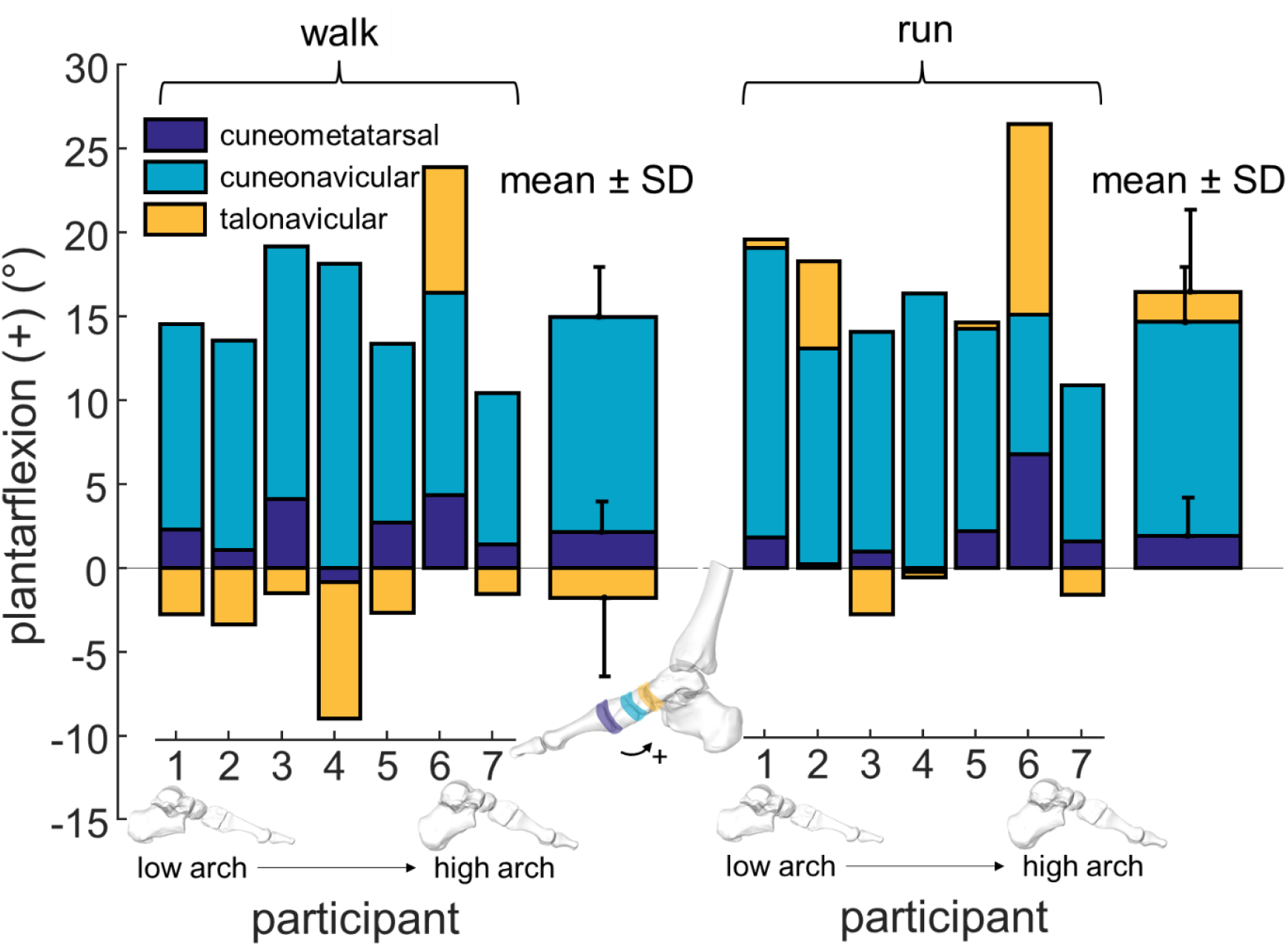
Contribution of the midfoot joints along the first ray to arch recoil (plantarflexion). Plantarflexion angle change between peak arch flattening and push-off is shown in bar form for each participant in both walking and running (navy/dark: cuneometatarsal; teal/medium: cuneonavicular; gold/light: talonavicular). Participants are ordered by unloaded arch plantarflexion angle. Coordinate systems are aligned with the first metatarsal at peak arch flattening (Supplementary Figure S1).

Enabling arch mobility has many important applications, including footwear design, understanding pathology, and surgical practice. The position of the rigid-arch tibia at push-off suggests that with an immobile arch, the body could lean forward to accommodate the tibial lean, modifying the posture of body segments and the muscular requirements for locomotion. This effect parallels the footwear literature as increasing the bending stiffness of the shoe’s sole can reduce arch deformation and recoil (Cigoja et al., 2020) and increase forward lean (Willwacher et al., 2014). In addition, when orthotic inserts restrict the arch’s motion, the metabolic cost of level-ground running is higher (Stearne et al., 2016). Overall, these studies suggest a relationship between arch mobility and forward lean, which may impact performance. Our results also have implications for people with naturally stiff feet or foot pathologies (such as osteoarthritis) that reduce mobility in the arch. When the tarsal joints are surgically fused, increased ankle power occurs in walking, supporting the idea that a rigid foot increases ankle plantarflexion velocity, force, or both (Beischer et al., 1999). Our method could also be used to predict dynamic motion patterns in surgical joint fusions. By mathematically fixing joints in known positions, we can elucidate potential changes along the kinematic chain. For example, we would expect that fusing the cuneonavicular joint would substantially impact propulsion, causing the foot to leave the ground early or increasing force requirements at the ankle. These results highlight the importance of preserving arch mobility in surgical practice and footwear design.

In conclusion, in bipedal walking and running, the mobility of the human medial arch does not directly propel the centre of mass. Instead, it works in tandem with the morphology of a high medial arch to facilitate upright locomotion through its effect on talus posture, ankle range of motion and ground contact time. We argue that while differences in medial arch height may visually distinguish hominins from other primates, our arch mobility is more critical to our ability to locomote on two feet. Thus, mapping morphology-mobility relationships in our extant relatives and humans, as well as forward-dynamic predictions of the fossil record, are necessary to understand our ancestors’ locomotory patterns.

## Materials and Methods

### BVR dataset

Seven young, physically active subjects (4F, 3M, mean±1SD, 23.3±3.0 years, 1.72±0.08m, 69.6±7.6kg, short IPAQ (Craig et al., 2003) moderate & vigorous physical activity: 477±325 min/week) were selected from a 20 participant pool to span the observed range of static arch heights. The selected participants walked and ran overground at a self-selected speed (walking (W) 1.6±0.1m/s, running (R) 3.0±0.4m/s) in flexible, thin-soled minimal shoes (7.5 mm sole, 0 mm heel-toe drop, Xero Prio Shoes, Broomfield, CO, USA) while biplanar videoradiography captured their foot bone motion (Skeletal Observation Lab, Queen’s University, Kingston, CAN). The experimental protocol was approved by Queen’s University Health Sciences and Affiliated Teaching Hospitals Research Ethics Board. All participants gave informed consent prior to participation in the data collection.

Three-dimensional positions and orientations of individual foot and ankle bones (tibia, calcaneus, talus, navicular, medial cuneiform, first metatarsal and first proximal phalanx) were measured using X-ray Reconstruction of Moving Morphology (Brainerd et al., 2010; Knörlein et al., 2016). This technology combines high-speed biplanar videoradiography ((W) 125Hz, (R) 250Hz) with bone models derived from a computed tomography (CT) scan to visualise rapid skeletal movement *in vivo*.

Three high-speed x-ray trials were collected for each participant (71kV, 125mA, shutter speed (W) 1250µs (R) 1000µs, resolution 2048 × 2048 pixels) as their right foot landed and pushed off in the x-ray capture volume. One trial was selected for analysis for x-ray image quality and appropriate participant foot placement. The biplanar videoradiography collection pipeline for the foot bones has been described previously (Kessler et al., 2019). Briefly, the high-speed cameras were calibrated using a custom calibration object, and the images were undistorted using open-source x-ray processing software (Knörlein et al., 2016) (XMALab, Providence, RI, USA).

A CT scan was taken of each participant’s right foot while supine, with a maximally plantarflexed ankle for improved in-plane resolution (Revolution HD; General Electric Medical Systems, Chicago, IL, USA; resolution: 0.317 mm x 0.317 mm x 0.625 mm). All bones (tibia, calcaneus, talus, navicular, medial cuneiform, first metatarsal, first proximal phalanx and first distal phalanx) were segmented (Mimics, Materialise, Leuven, Belgium). Tessellated meshes depicted the bone surfaces and were used to establish inertial coordinate systems. The coordinate systems’ origin was located at the centroid of each bone, and the three axes aligned with the principal directions of the moment of inertia tensor (Eberly et al., 1991). The axes were re-labelled such that the x-axis was lateral, with positive angles about this axis indicating dorsiflexion. Specialised coordinate systems with a cylinder fitted to the talar and tibia domes were used to measure ankle dorsi/plantarflexion.

Partial volumes generated from the bone masks formed digitally reconstructed radiographs (Miranda et al., 2011). Custom software (Autoscoper, Brown University, Providence, RI, USA) semi-manually measured the orientation and translation of the bones of interest using the digitally reconstructed radiographs and undistorted x-ray images. The digitally reconstructed radiographs were manually aligned with the two x-ray views, and a particle swarm algorithm optimised the normalised cross-correlation values (Akhbari et al., 2021). A 3D visualisation of the bone positions ensured that no collisions occurred between adjacent bones (Welte et al., 2022).

### Angles

Dorsiflexion (+) /plantarflexion (-) is measured as the Tait-Bryan angle of the distal bone relative to the proximal bone using a YZX sequence to prioritise x-axis dorsi/plantarflexion. MTPJ dorsiflexion measures the first proximal phalanx’s motion relative to the first metatarsal. Arch plantarflexion and dorsiflexion refer to the first metatarsal’s sagittal motion relative to the calcaneus and are referred to as arch angle. Arch angle is described as flattening (dorsiflexion) in early- and mid-stance, and as recoil (plantarflexion) in propulsion. Ankle dorsi/plantarflexion measures the talus’s motion relative to the tibia. Range of motion for ankle plantarflexion and arch recoil is measured between peak arch flattening and peak MTPJ dorsiflexion.

To measure the contributions of the medial column to propulsion, we measured the plantarflexion of each arch bone-pair (talonavicular, cuneonavicular, cuneometatarsal) using the orientation of the inertial coordinate system of the first metatarsal, aligned at peak arch flattening (Extended Data Figure 1). The convention for this analysis was to report plantarflexion as positive by switching the sign of the Tait-Bryan angle. The range of motion for each joint was again measured between peak arch flattening and peak MTPJ dorsiflexion.

Tibial lean was measured as the global orientation of the tibia relative to the global axes. The X angle of a Tait-Bryan YZX sequence of the tibial inertial coordinate system relative to the global axis measured the tibial lean. The tibial anatomical coordinate system was defined specifically for this measurement, with the z-axis aligned with a cylinder fitted to the long axis of the tibial shaft. The x-axis is directed laterally (approximately intersecting the medial malleolus), and the y-axis is the anteriorly directed mutual perpendicular to the x- and z-axes.

### Arch Height

Arch height was measured in a static seated position. The participants were instructed to place their right barefoot in front of their left foot in the capture volume and to distribute their weight evenly between their legs. The position and orientation of the arch bones and first proximal phalanx were measured using the previously described methods for processing biplanar x-ray data.

A slightly modified arch height index (AHI) contextualised the range of foot types (Butler et al., 2008). AHI is typically measured with a specialised device and is the quotient of the arch height at 50% of the total foot length and the truncated foot length. Here, the medial arch was oriented by the principal component axes of the arch bone vertices. The first principal component represented the anterior direction of the foot and the second principal component represented the height of the arch. The truncated foot length measured the distance from the most posterior point of the heel to the anterior tip of the first metatarsal. As we could not measure the pose of the longest distal phalanx, we fixed the first distal phalanx with the first proximal phalanx. Foot length was measured from the previously calculated posterior heel point to the tip of the first distal phalanx. The highest vertex of the arch bones within 1 mm of the 50% length of the foot was selected as the dorsal arch height. The dorsal arch height was then divided by the truncated foot length to measure AHI.

Arch angle was also measured in the static seated position. AHI and arch angle were linearly correlated (p < 0.01, R^2^= 0.87); thus, we selected arch static dorsiflexion angle for our analyses as it was more similar to our dynamic measures.

### Rigid Foot Model

We tested the contributions of a rigid arch to propulsion by mathematically locking the arch bones with the first metatarsal at the beginning of arch recoil (see Supplementary Methods 1), which is the flattest arch position during stance. The arch posture (i.e. fully flat or fully recoiled) did not change our outcomes. The arch bones, with no relative motion between them, were driven with the motion of the first metatarsal through propulsion. The end of propulsion was defined to be the maximum MTPJ dorsiflexion angle. Relative tibiotalar motion remained consistent between rigid and moving arch propulsion. Comparisons were made between the modelled rigid arch and the measured naturally recoiling arch for each participant. All modelling and optimisation described in this section was conducted in MATLAB R2020b (Mathworks, Natick, MA, USA).

To test the contributions of the foot’s levering motion (rigid) and arch recoil (moving) to centre of mass propulsion, we measured the position of the talar centroid for the rigid and recoiling arch. The position of the talar centroid is a proxy for the ankle’s ability to lift the centre of mass. The talar centroid was projected into the inertial coordinate system of the first proximal phalanx to standardise the direction of take-off among participants. We measured the height and forward progression of the talar centroid in three analyses: first, at the end of propulsion, with the same contact time and therefore levering motion between rigid and moving arches; second, when the rigid arch’s tibia aligned with the moving arch’s tibia at push-off; and third, with both levering motion and push-off tibia position maintained. In the second analysis, tibia alignment was measured as the maximum value of the dot product of the vector aligned with the tibial shaft in each condition. In the third analysis, the tibia was rotated about its helical axis by the angle between the global positions of the rigid and moving-arch tibiae. The translation of the newly rotated tibia was optimised such that there were no bone collisions while minimising the mean distance between the tibia and talus.

A two-tailed Wilcoxon signed-ranks test measured the difference in talar height between the rigid and moving arches for walking and running as Shapiro-Wilks normality tests indicated that the distributions were not normal. Significance was set at α = 0.05. Effect sizes are reported as the difference between the proportions of favourable and unfavourable outcomes, with 0 indicating no effect and 1 indicating all pairs behaved the same way (Kerby, 2014). All reported values are mean ± one standard deviation unless otherwise indicated.

### Multiple Regression Model

The influence of arch mobility and arch height on ankle kinematic measurements was analysed using two multiple regression models. The predictor variables were arch recoil range of motion, static arch angle, and a categorical variable indicating whether the trial was a walk or a run. The response variables were the magnitude of talar displacement in the anterior and superior directions (from peak arch flattening to peak MTPJ dorsiflexion) and ankle plantarflexion range of motion. Assumptions of variable collinearity and homoscedasticity, as well as independence and normality of residual values, were met. Significance was set at α = 0.05. Statistical analysis was completed in MATLAB R2020b (Mathworks, Natick, MA, USA), using the fitlm function.

### Chimpanzee dataset

Chimpanzee data were collected previously from three subadult male chimpanzee subjects (age: 5.5 ± 0.2 yrs; 26.5 ± 6.7 kg) (O’Neill et al., 2015). Chimpanzees were housed at an Association for Assessment and Accreditation of Animal Laboratory Care International-approved facility, and all experimental protocols were approved by Stony Brook University’s Institutional Animal Care and Use Committee. Chimpanzees were previously trained to walk bipedally using positive reinforcement. During data collection, chimpanzees were encouraged to walk bipedally by an animal trainer who used food and juice rewards. Four high-speed video cameras recording at 150 Hz were used to capture motion as the subject walked on a flat 11-meter runway at self-selected speeds. We analysed a total of 13 bipedal steps (Subject A: 5 steps; Subject B: 1 step; Subject C: 7 steps), and subjects walked at an average speed of 1.13 ± 0.11 m/s (0.35 ± 0.09 Froude). Tibial lean was calculated as the angle between the global vertical axis and the vector connecting markers that were applied to the lateral malleolus and the fibular head.

## Acknowledgments

Funding for this work came from an Ontario Early Researcher Award, an NSERC Discovery Grant (RGPIN/04688-2015), an NSERC’s Postgraduate Scholarship – Doctoral and the Pedorthic Research Foundation of Canada. Shoes were provided by Xero Shoes.

## Supplementary Methods 1

Here, we provide the math that rigidly locks the arch, and the cases demonstrated in *Mobility of the human foot’s medial arch enables upright bipedal locomotion*.

### Conventions

[T] is the 4×4 matrix that moves a rigid body from computed tomography (CT) space to x-ray (xr) space. T is composed of a 3×3 rotation matrix and 3×1 translation matrix.

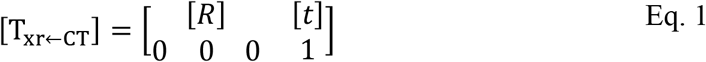

The points on a bone can be moved using Eq. 1:

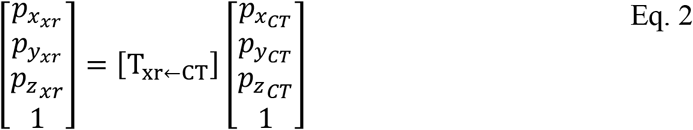

The transform for a specific bone (i.e. mt1 = first metatarsal) is denoted in superscript of the transform. The frame (i) is given outside the brackets (peak arch flattening = pa).

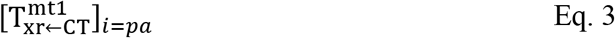

All output transforms given here transform the bone’s vertices in CT space to x-ray space.

#### Case 1: Fix the arch bones rigidly with the first metatarsal at peak arch flattening

**Lock the arch**. To fix the talus (tal) at any frame (i) with the first metatarsal (mt1) at peak arch flattening (i=pa):

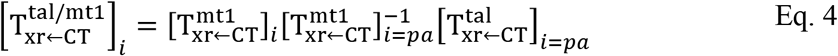

This locks the position of the talus relative to the first metatarsal at the frame pa, and then moves it with the first metatarsal’s motion.

#### Case 2: Maintain some joints’ natural motion, but still lock the arch with the first metatarsal

**Lock the arch, but maintain tibio-talar motion**. The arch is locked to the first metatarsal below the tibiotalar joint. To maintain the relative motion of the tibia (tib) relative to a talus fixed with the first metatarsal:

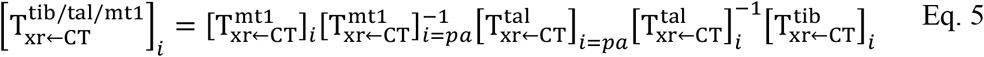

Note that the first three transforms can simplify to Eq. 4.

#### Case 3: Fix a joint to the position at a different time than when the arch is locked

**Lock the arch at peak arch flattening, but keep the tibio-talar joint in the take-off (to) position**.

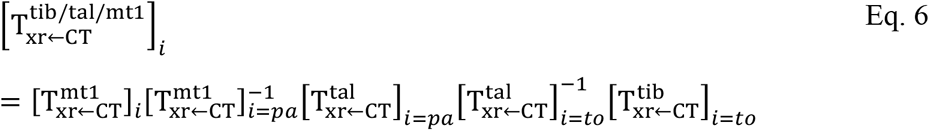

Again, note the first three transforms are the same as Eq. 4. The only difference between Eq 5 and 6 are the frame at which the last two transforms are taken.

**Fig. S1.**
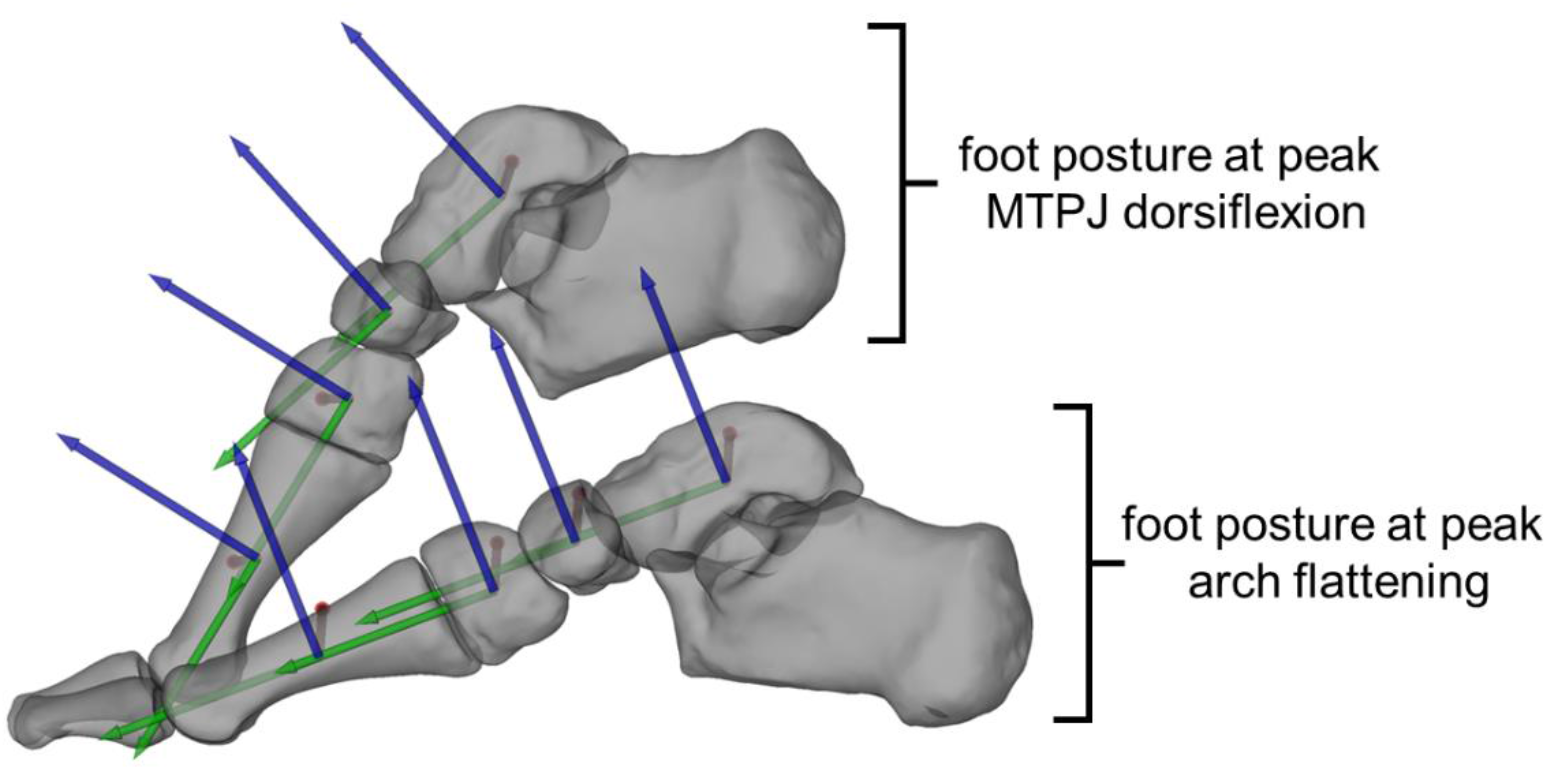
Medial arch co-ordinate systems. Co-ordinate systems of the arch bones, fixed to align with the orientation of the inertial co-ordinate system of the first metatarsal at peak arch flattening. A representative participant’s foot posture is shown during a run at peak arch flattening and peak metatarsophalangeal joint (MTPJ) dorsiflexion.

**Movie S1 (separate file)**. Motion of the recoiling and modelled rigid medial arch. The levering motion of the first metatarsal is included on the right, but has been removed to show the relative motion of the recoiling arch to the rigid arch on the left.

## Notes

https://www.dropbox.com/sh/5tk3oxqmqqnd5st/AAD4ETo2dkmZwCwwZTEJLicra?dl=0

### Competing Interest Statement

The authors have declared no competing interest.

## References

Akhbari B, Morton AM, Moore DC, Crisco JJ. 2021. Biplanar Videoradiography to Study the Wrist and Distal Radioulnar Joints. JoVE (Journal of Visualized Experiments) e62102. doi:10.3791/62102

Akhbari B, Morton AM, Moore DC, Weiss A-PC, Wolfe SW, Crisco JJ. 2019. Accuracy of biplane videoradiography for quantifying dynamic wrist kinematics. Journal of Biomechanics 92:120–125. doi:10.1016/j.jbiomech.2019.05.040

Arndt A, Wolf P, Liu A, Nester C, Stacoff A, Jones R, Lundgren P, Lundberg A. 2007. Intrinsic foot kinematics measured in vivo during the stance phase of slow running. Journal of Biomechanics 40:2672–2678. doi:10.1016/j.jbiomech.2006.12.009

Bates KT, Collins D, Savage R, McClymont J, Webster E, Pataky TC, D’Aout K, Sellers WI, Bennett MR, Crompton RH. 2013. The evolution of compliance in the human lateral mid-foot. Proc Biol Sci 280. doi:10.1098/rspb.2013.1818

Beischer AD, Brodsky JW, Polio FE, Peereboom J. 1999. Functional Outcome and Gait Analysis After Triple or Double Arthrodesis. Foot Ankle Int 20:545–553. doi:10.1177/107110079902000902

Bennett MB, Ker RF, Alexander RM. 1989. Elastic strain energy storage in the feet of running monkeys. Journal of Zoology 217:469–475. doi:10.1111/j.1469-7998.1989.tb02502.x

Bojsen-Møller F. 1979. Calcaneocuboid joint and stability of the longitudinal arch of the foot at high and low gear push off. J Anat 129:165–176.

Brainerd EL, Baier DB, Gatesy SM, Hedrick TL, Metzger KA, Gilbert SL, Crisco JJ. 2010. X-ray reconstruction of moving morphology (XROMM): precision, accuracy and applications in comparative biomechanics research. J Exp Zool A Ecol Genet Physiol 313:262–279. doi:10.1002/jez.589

Bramble DM, Lieberman DE. 2004. Endurance running and the evolution of Homo. Nature 432:345–352. doi:10.1038/nature03052

Butler RJ, Hillstrom H, Song J, Richards CJ, Davis IS. 2008. Arch Height Index Measurement System: Establishment of Reliability and Normative Values. Journal of the American Podiatric Medical Association 98:102–106. doi:10.7547/0980102

Carrier DR, Heglund NC, Earls KD. 1994. Variable gearing during locomotion in the human musculoskeletal system. Science 265:651–653.

Cavagna GA, Heglund NC, Taylor CR. 1977. Mechanical work in terrestrial locomotion: two basic mechanisms for minimizing energy expenditure. American Journal of Physiology-Regulatory, Integrative and Comparative Physiology 233:R243–R261. doi:10.1152/ajpregu.1977.233.5.R243

Cigoja S, Asmussen MJ, Firminger CR, Fletcher JR, Edwards WB, Nigg BM. 2020. The Effects of Increased Midsole Bending Stiffness of Sport Shoes on Muscle-Tendon Unit Shortening and Shortening Velocity: a Randomised Crossover Trial in Recreational Male Runners. Sports Medicine - Open 6:9. doi:10.1186/s40798-020-0241-9

Cornwall MW, McPoil TG. 2011. Relationship between static foot posture and foot mobility. J Foot Ankle Res 4:4. doi:10.1186/1757-1146-4-4

Craig CL, Marshall AL, Sj??Str??M M, Bauman AE, Booth ML, Ainsworth BE, Pratt M, Ekelund U, Yngve A, Sallis JF, Oja P. 2003. International Physical Activity Questionnaire: 12-Country Reliability and Validity: Medicine & Science in Sports & Exercise 35:1381–1395. doi:10.1249/01.MSS.0000078924.61453.FB

D’Août K, Aerts P, Clercq DD, Meester KD, Elsacker LV. 2002. Segment and joint angles of hind limb during bipedal and quadrupedal walking of the bonobo (Pan paniscus). American Journal of Physical Anthropology 119:37–51. doi:10.1002/ajpa.10112

DeSilva J, McNutt E, Benoit J, Zipfel B. 2019. One small step: A review of Plio-Pleistocene hominin foot evolution. American Journal of Physical Anthropology 168:63–140. doi:10.1002/ajpa.23750

DeSilva JM. 2010. Revisiting the “midtarsal break.” American Journal of Physical Anthropology 141:245–258. doi:10.1002/ajpa.21140

DeSilva JM, Bonne-Annee R, Swanson Z, Gill CM, Sobel M, Uy J, Gill SV. 2015. Midtarsal break variation in modern humans: Functional causes, skeletal correlates, and paleontological implications. American Journal of Physical Anthropology 156:543–552. doi:10.1002/ajpa.22699

Eberly D, Lancaster J, Alyassin A. 1991. On gray scale image measurements: II. Surface area and volume. CVGIP: Graphical Models and Image Processing 53:550–562. doi:10.1016/1049-9652(91)90005-5

Elftman H., Manter J. 1935. The Evolution of the Human Foot, with Especial Reference to the Joints. J Anat 70:56–67.

Elftman Herbert, Manter J. 1935. Chimpanzee and human feet in bipedal walking. American Journal of Physical Anthropology 20:69–79. doi:10.1002/ajpa.1330200109

Farris DJ, Kelly LA, Cresswell AG, Lichtwark GA. 2019. The functional importance of human foot muscles for bipedal locomotion. PNAS 116:1645–1650. doi:10.1073/pnas.1812820116

Farris DJ, Sawicki GS. 2012. The mechanics and energetics of human walking and running: a joint level perspective. Journal of The Royal Society Interface 9:110–118. doi:10.1098/rsif.2011.0182

Greiner TM, Ball KA. 2014. Kinematics of primate midfoot flexibility. American Journal of Physical Anthropology 155:610–620. doi:10.1002/ajpa.22617

Griffin NL, D’Août K, Richmond B, Gordon A, Aerts P. 2010. Comparative in vivo forefoot kinematics of Homo sapiens and Pan paniscus. Journal of Human Evolution 59:608–619. doi:10.1016/j.jhevol.2010.07.017

Hicks JH. 1955. The foot as a support. CTO 25:34–45. doi:10.1159/000141055

Hicks JH. 1954. The mechanics of the foot. II. The plantar aponeurosis and the arch. J Anat 88:25–30.

Holowka NB, Lieberman DE. 2018. Rethinking the evolution of the human foot: insights from experimental research. Journal of Experimental Biology 221:jeb174425. doi:10.1242/jeb.174425

Holowka NB, O’Neill MC, Thompson NE, Demes B. 2017a. Chimpanzee and human midfoot motion during bipedal walking and the evolution of the longitudinal arch of the foot. Journal of Human Evolution 104:23–31. doi:10.1016/j.jhevol.2016.12.002

Holowka NB, O’Neill MC, Thompson NE, Demes B. 2017b. Chimpanzee ankle and foot joint kinematics: Arboreal versus terrestrial locomotion. American Journal of Physical Anthropology 164:131–147. doi:10.1002/ajpa.23262

Kelly LA, Lichtwark G, Cresswell AG. 2014. Active regulation of longitudinal arch compression and recoil during walking and running. Journal of The Royal Society Interface 12:20141076–20141076. doi:10.1098/rsif.2014.1076

Ker RF, Bennett MB, Bibby SR, Kester RC, Alexander RMcN. 1987. The spring in the arch of the human foot. Nature 325:147–149. doi:10.1038/325147a0

Kerby DS. 2014. The Simple Difference Formula: An Approach to Teaching Nonparametric Correlation. Comprehensive Psychology 3:11.IT.3.1. doi:10.2466/11.IT.3.1

Kessler SE, Rainbow MJ, Lichtwark GA, Cresswell AG, D’Andrea SE, D’Andrea SE, D’Andrea SE, Konow N, Kelly LA. 2019. A Direct Comparison of Biplanar Videoradiography and Optical Motion Capture for Foot and Ankle Kinematics. Frontiers in Bioengineering and Biotechnology 7. doi:10.3389/fbioe.2019.00199

Knörlein BJ, Baier DB, Gatesy SM, Laurence-Chasen JD, Brainerd EL. 2016. Validation of XMALab software for marker-based XROMM. Journal of Experimental Biology 219:3701–3711. doi:10.1242/jeb.145383

Kondo M, Iwamoto Y, Kito N. 2021. Relationship between forward propulsion and foot motion during gait in healthy young adults. Journal of Biomechanics 121:110431. doi:10.1016/j.jbiomech.2021.110431

Lovejoy CO, Latimer B, Suwa G, Asfaw B, White TD. 2009. Combining Prehension and Propulsion: The Foot of Ardipithecus ramidus. Science 326:72–72e8. doi:10.1126/science.1175832

McNutt EJ, Zipfel B, DeSilva JM. 2018. The evolution of the human foot. Evol Anthropol 27:197–217. doi:10.1002/evan.21713

Miranda DL, Schwartz JB, Loomis AC, Brainerd EL, Fleming BC, Crisco JJ. 2011. Static and Dynamic Error of a Biplanar Videoradiography System Using Marker-Based and Markerless Tracking Techniques. Journal of Biomechanical Engineering 133:121002. doi:10.1115/1.4005471

Morton DJ. 1924. EVOLUTION OF THE LONGITUDINAL ARCH OF THE HUMAN FOOT. JBJS 6:56–90.

Negishi T, Ito K, Hosoda K, Nagura T, Ota T, Imanishi N, Jinzaki M, Oishi M, Ogihara N. 2021. Comparative radiographic analysis of three-dimensional innate mobility of the foot bones under axial loading of humans and African great apes. Royal Society Open Science 8:211344. doi:10.1098/rsos.211344

Neptune RR, Kautz SA, Zajac FE. 2001. Contributions of the individual ankle plantar flexors to support, forward progression and swing initiation during walking. Journal of Biomechanics 34:1387–1398. doi:10.1016/S0021-9290(01)00105-1

O’Neill MC, Lee L-F, Demes B, Thompson NE, Larson SG, Stern JT, Umberger BR. 2015. Three-dimensional kinematics of the pelvis and hind limbs in chimpanzee (Pan troglodytes) and human bipedal walking. Journal of Human Evolution 86:32–42. doi:10.1016/j.jhevol.2015.05.012

Pontzer H. 2017. Economy and Endurance in Human Evolution. Current Biology 27:R613–R621. doi:10.1016/j.cub.2017.05.031

Sarrafian SK. 1987. Functional Characteristics of the Foot and Plantar Aponeurosis under Tibiotalar Loading. Foot Ankle Int 8:4–18. doi:10.1177/107110078700800103

Sichting F, Holowka NB, Ebrecht F, Lieberman DE. 2020. Evolutionary anatomy of the plantar aponeurosis in primates, including humans. Journal of Anatomy 00:1–21. doi:10.1111/joa.13173

Sockol MD, Raichlen DA, Pontzer H. 2007. Chimpanzee locomotor energetics and the origin of human bipedalism. Proceedings of the National Academy of Sciences 104:12265–12269. doi:10.1073/pnas.0703267104

Stearne SM, McDonald KA, Alderson JA, North I, Oxnard CE, Rubenson J. 2016. The Foot’s Arch and the Energetics of Human Locomotion. Scientific Reports 6. doi:10.1038/srep19403

Susman RL. 1983. Evolution of the human foot: evidence from Plio-Pleistocene hominids. Foot Ankle 3:365–376.

Venkadesan M, Yawar A, Eng CM, Dias MA, Singh DK, Tommasini SM, Haims AH, Bandi MM, Mandre S. 2020. Stiffness of the human foot and evolution of the transverse arch. Nature 579:97–100. doi:10.1038/s41586-020-2053-y

Vereecke E, D’Août K, Clercq DD, Elsacker LV, Aerts P. 2003. Dynamic plantar pressure distribution during terrestrial locomotion of bonobos (Pan paniscus). American Journal of Physical Anthropology 120:373–383. doi:10.1002/ajpa.10163

Vereecke EE, Aerts P. 2008. The mechanics of the gibbon foot and its potential for elastic energy storage during bipedalism. Journal of Experimental Biology 211:3661–3670. doi:10.1242/jeb.018754

Waters RL, Mulroy S. 1999. The energy expenditure of normal and pathologic gait. Gait Posture 9:207–231. doi:10.1016/s0966-6362(99)00009-0

Welte L, Dickinson A, Arndt A, Rainbow MJ. 2022. Biplanar Videoradiography Dataset for Model-based Pose Estimation Development and New User Training. JoVE (Journal of Visualized Experiments) e63535. doi:10.3791/63535

Willwacher S, König M, Braunstein B, Goldmann J-P, Brüggemann G-P. 2014. The gearing function of running shoe longitudinal bending stiffness. Gait & Posture 40:386–390. doi:10.1016/j.gaitpost.2014.05.005

Zifchock RA, Davis I, Hillstrom H, Song J. 2006. The Effect of Gender, Age, and Lateral Dominance on Arch Height and Arch Stiffness. Foot & Ankle International 27:367–372. doi:10.1177/107110070602700509

